# CyMIRA: The Cytonuclear Molecular Interactions Reference for *Arabidopsis*

**DOI:** 10.1101/614487

**Authors:** Evan S. Forsythe, Joel Sharbrough, Justin C. Havird, Jessica M. Warren, Daniel B. Sloan

## Abstract

The function and evolution of eukaryotic cells depends upon direct molecular interactions between gene products encoded in nuclear and cytoplasmic genomes. Understanding how these cytonuclear interactions drive molecular evolution and generate genetic incompatibilities between isolated populations and species is of central importance to eukaryotic biology. Plants are an outstanding system to investigate such effects because of their two different genomic compartments present in the cytoplasm (mitochondria and plastids) and the extensive resources detailing subcellular targeting of nuclear-encoded proteins. However, the field lacks a consistent classification scheme for mitochondrial- and plastid-targeted proteins based on their molecular interactions with cytoplasmic genomes and gene products, which hinders efforts to standardize and compare results across studies. Here, we take advantage of detailed knowledge about the model angiosperm *Arabidopsis thaliana* to provide a curated database of plant cytonuclear interactions at the molecular level. CyMIRA (<u>Cy</u>tonuclear <u>M</u>olecular <u>I</u>nteractions <u>R</u>eference for *<u>A</u>rabidopsis*) is available at http://cymira.colostate.edu/ and https://github.com/dbsloan/cymira and will serve as a resource to aid researchers in partitioning evolutionary genomic data into functional gene classes based on organelle targeting and direct molecular interaction with cytoplasmic genomes and gene products. It includes 11 categories (and 27 subcategories) of different cytonuclear complexes and types of molecular interactions, and it reports residue-level information for cytonuclear contact sites. We hope that this framework will make it easier to standardize, interpret and compare studies testing the functional and evolutionary consequences of cytonuclear interactions.

## INTRODUCTION

The endosymbiotic history of eukaryotes has resulted in cells that are operated under divided genetic control between nuclear and cytoplasmic (i.e., mitochondrial and plastid) genomes. Core eukaryotic functions depend on integration and coevolution between these genomic compartments. The level of integration extends down to direct molecular interactions within multisubunit enzyme complexes (Rand, et al. 2004). For example, the major enzymes in mitochondria and plastids such as oxidative phosphorylation (OXPHOS) complexes, photosynthetic machinery, and ribosomes are ‘chimeric’ in the sense that they are composed of gene products from both nuclear and cytoplasmic genomes. This organization reflects an evolutionary history in which many of the genes ancestrally present in cytoplasmic genomes have been replaced by a combination of gene transfer to the nucleus and substitution by existing nuclear genes (Sloan, et al. 2018). There are also extensive interactions between cytoplasmic RNAs and nuclear-encoded proteins that are responsible for post-transcriptional processes, such as transcript end-processing, intron splicing, RNA editing, base modifications, and tRNA aminoacylation (Germain, et al. 2013; Salinas-Giegé, et al. 2015). Furthermore, many nuclear-encoded proteins must directly interact with the cytoplasmic genomes themselves to mediate processes of DNA replication, repair, recombination, and transcription (Zhang, et al. 2016; Gualberto and Newton 2017).

The intimacy of these interactions has made them an attractive arena for studying molecular coevolution, especially because they can elucidate the consequences of genes evolving in very different genomic contexts (e.g., differences in mutation rates, replication and expression mechanisms, frequency of recombination, effective population sizes, and modes of inheritance). Not surprisingly, disruption of cytonuclear interactions can have significant functional consequences, and genetic incompatibilities between nuclear and cytoplasmic genomes contribute to reproductive isolation in many systems (Burton, et al. 2013; Hill 2015; Sloan, et al. 2017). It is possible that cytonuclear incompatibilities evolve at a faster pace than nuclear-nuclear incompatibilities because of the differences in genome evolution and the conflicting genealogical histories that can often distinguish these compartments (Burton and Barreto 2012; Toews and Brelsford 2012).

To test such hypotheses, it is often useful to compare nuclear-encoded proteins that are involved in direct cytonuclear molecular interactions against relevant ‘control’ proteins. For example, classic studies in animals have taken advantage of OXPHOS complex II (succinate dehydrogenase), which is entirely nuclear-encoded, in order to make comparisons with the other OXPHOS complexes, which are all chimeric (Ellison and Burton 2006). In the current genomic era, it has become increasingly popular for evolutionary studies to partition nuclear gene content into categories based on whether they are targeted to mitochondria/plastids and whether they are involved in direct molecular interactions with cytoplasmic genomes and gene products within these organelles (Barreto and Burton 2013; Rogell, et al. 2014; Pett and Lavrov 2015; Sloan, et al. 2015; Zhang, et al. 2015; Adrion, et al. 2016; Rockenbach, et al. 2016; Weng, et al. 2016; Zhang, et al. 2016; Eslamieh, et al. 2017; Havird, et al. 2017; Sharbrough, et al. 2017; Barreto, et al. 2018; Forsythe, et al. 2018; Morales, et al. 2018; Ferreira, et al. 2019; Li, et al. 2019; Yan, et al. 2019; Zaidi and Makova 2019). Such approaches are an effective means to investigate the evolutionary effect of organelle targeting and molecular interactions. Because plants contain two endosymbiotically derived organelles, they are an especially appealing system in which to study such questions. However, comparing across studies can be challenging because of the variable ways in which authors classify and partition gene sets. Although there are many excellent databases with gene-specific information on subcellular targeting in plants (Table 1), none of these provide comprehensive information about direct cytonuclear interactions at the level of protein subunits and amino-acid residues. To address this limitation, we have taken advantage of the extensive work on cytonuclear biology in the model angiosperm *Arabidopsis thaliana* to create the Cytonuclear Molecular Interactions Reference for *Arabidopsis* (CyMIRA).

**Table 1.** Set of nine databases with information on subcellular localization of proteins in plants that were used for automated curations of targeting predictions. Counts reflect number of genes in each targeting category.

| Database | Targeting Predictions |  |  | Reference |
| --- | --- | --- | --- | --- |
|  | Mito | Plastid | Dual |  |
| SUBA predicted | 2370 | 2644 | 97 | (Hooper, et al. 2017) |
| eSLDB | 848 | 4427 | 69 | (Pierleoni, et al. 2007) |
| PA-GOSUB | 985 | 730 | 14 | (Lu, et al. 2005) |
| SUBA experimental | 1217 | 2128 | 785 | (Hooper, et al. 2017) |
| SWISS PROT | 311 | 657 | 20 | (Boutet, et al. 2007) |
| TAIR | 397 | 1598 | 266 | (Reiser, et al. 2017) |
| LocDB | 446 | 1527 | 234 | (Rastogi and Rost 2011) |
| PPDB | 327 | 1570 | 73 | (Sun, et al. 2009) |
| Organelle DB | 512 | 276 | 11 | (Wiwatwattana, et al. 2007) |

## MATERIALS AND METHODS

### Curation of mitochondrial and plastid targeting databases

To identify mitochondrial- and plastid-targeted genes, we integrated predictions from nine existing databases (Table 1). Based on these datasets, we classified all nuclear-encoded proteins in the *A. thaliana* Araport11 genome annotation into five targeting categories: mitochondrial, plastid, dual (both mitochondrial and plastid), other, or unknown. Because of the cytonuclear focus of this project, in cases where organelle-targeted proteins were known to have additional subcellular localizations, we still classified them based on their mitochondrial/plastid targeting status alone. To classify a protein as having an organellar localization, we required it to be identified as such in at least two different databases. Because it is well documented that many plant proteins play a dual functional role in both the mitochondria and plastids (Carrie and Small 2013), we assigned genes to the dual-targeted category as long as there were at least two databases supporting targeting to the mitochondria and at least two supporting targeting to the plastids. It was possible (although not required) for these to be the same two databases because the selected databases explicitly classify some genes as dual targeted. Some of these automated database classifications were subsequently refined based on manual curation of direct molecular interactions as described below.

### Curation of direct cytonuclear molecular interaction

We conducted a literature-based curation to generate a resource that could distinguish nuclear proteins that are simply targeted to mitochondria and plastids from those that are involved in direct and intimate interactions with cytoplasmic genomes or their gene products. We assigned genes to 11 types of cytonuclear enzyme complexes and molecular interactions, which are further divided into 27 subcategories (Table 2).

**Table 2.** List of functional categories used in manual curation of direct cytonuclear interactions. Counts reflect number of genes in each targeting category. Key references are listed by category. More extensive literature references are provided in Table S1.

| Category | Subcategory | Mito | Plastid | Dual | Key Reference(s) |
| --- | --- | --- | --- | --- | --- |
| **ACCase** |  | **0** | **4** | **0** | (Sasaki and Nagano 2004) |
| **Chlororibosome** |  | **0** | **42** | **0** | (Bonen and Calixte 2006; Sloan, et al. 2014) |
|  | Large subunit | 0 | 31 | 0 | (Bieri, et al. 2017) |
|  | Small subunit | 0 | 11 | 0 | (Tiller, et al. 2012) |
| **Clp protease** |  | **0** | **15** | **0** | (Nishimura and van Wijk 2015) |
| **DNA-RRR** |  | **11** | **8** | **17** | (Zhang, et al. 2016; Gualberto and Newton 2017) |
| **TAT complex** |  | **1** | **0** | **0** | (Carrie, et al. 2016) |
| **Mitoribosome** |  | **88** | **0** | **0** | (Waltz, et al. 2019) |
|  | Large subunit | 41 | 0 | 0 |  |
|  | Small subunit | 47 | 0 | 0 |  |
| **OXPHOS** |  | **91** | **0** | **0** | (Senkler, et al. 2017) |
|  | Complex I | 48 | 0 | 0 |  |
|  | Complex III | 14 | 0 | 0 |  |
|  | Complex IV | 14 | 0 | 0 |  |
|  | Complex V | 15 | 0 | 0 |  |
| **Photosynthesis** |  | **0** | **67** | **0** |  |
|  | ATP synthase | 0 | 3 | 0 | (Friso, et al. 2004) |
|  | Cytochrome b6f | 0 | 2 | 0 | (Friso, et al. 2004) |
|  | NDH | 0 | 18 | 0 | (Shikanai 2016) |
|  | PSI | 0 | 18 | 0 | (Jensen, et al. 2007) |
|  | PSII | 0 | 22 | 0 | (van Bezouwen, et al. 2017) |
|  | Rubisco | 0 | 4 | 0 | (Izumi, et al. 2012) |
| **PPR** |  | **308** | **110** | **36** | (Cheng, et al. 2016) |
| **Transcription and transcript maturation** |  | **33** | **46** | **5** |  |
|  | Intron splicing | 7 | 7 | 1 | (de Longevialle, et al. 2010) |
|  | mTERF | 17 | 11 | 0 | (Shevtsov, et al. 2018) |
|  | RNA polymerase | 1 | 1 | 1 | (Kühn, et al. 2007) |
|  | rRNA base modification | 1 | 2 | 0 | (Yu, et al. 2008) |
|  | Sigma factor | 0 | 6 | 0 | (Zhang, et al. 2015) |
|  | Transcript end processing | 5 | 5 | 3 | (Perrin, et al. 2004; Stoll and Binder 2016) |
|  | tRNA base modification | 2 | 14 | 0 | (Chen, et al. 2010) |
| **tRNA aminoacylation** |  | **3** | **1** | **24** | (Duchêne, et al. 2005) |
| **Total** |  | **535** | **293** | **82** |  |

Because of the manual nature of this curation, our classifications often required judgment calls and special considerations. With respect to major multi-subunit enzymes, we aimed to restrict our classification to the core complex, excluding proteins such as assembly factors involved in more transient interactions (e.g., Lu 2016; Ligas, et al. 2019).

One of the largest classes of genes involved in plant cytonuclear interactions is the RNA-binding pentatricopeptide repeat (PPR) family (Schmitz-Linneweber and Small 2008). These proteins are overwhelmingly targeted to the mitochondria and plastids where they play diverse roles in RNA processing and maturation. We classified six specialized PPRs as components of the mitochondrial ribosome (Waltz, et al. 2019) or as functioning in tRNA end processing (Gobert, et al. 2010). The remaining PPRs were assigned to their own category. Even though many PPRs still lack detailed functional characterization, we considered these examples of direct cytonuclear interactions because of their near universal role in binding cytoplasmic transcripts. A total of 109 PPRs (24%) were not identified as mitochondrial or plastid targeted based on our automated database curation. In these cases, we reassigned their targeting classification using The Arabidopsis Information Resource (TAIR) Gene Ontology (GO) cellular component designations (Berardini, et al. 2015). As a result, all PPRs were assigned as mitochondrial and/or plastid targeted, with the exception of only nine genes (AT1G06150, AT1G77150, AT2G20720, AT3G13150, AT3G47530, AT3G58590, AT5G09320, AT5G15300, AT5G44230), which we excluded from the direct-interaction dataset. A large portion of PPR genes function as specificity factors in C-to-U RNA editing of organellar transcripts. Therefore, RNA editing interactions are effectively subsumed within the PPR category. Although other types of nuclear proteins have been found to function in RNA editing (Sun, et al. 2016), we are not aware of any evidence that these directly bind to organellar transcripts, so they were not classified as directly interacting.

Mitochondrial transcription termination factors (mTERFs) are another sizeable family of organelle-targeted nucleic-acid binding proteins (Shevtsov, et al. 2018). Similar to how we handled PPRs, we defined mTERFs as their own subcategory within the transcription and transcript maturation category, even though many individual mTERF genes await functional characterization.

Although our manual curation of direct cytonuclear interactions overwhelmingly agreed with general subcellular targeting predictions from our database summary, there were 189 genes (21%, including 100 PPRs; see above) for which the automated targeting predictions did not include the organelle(s) found in our manual analysis. In such cases, we updated the original automated targeting call by adding the location of direct cytonuclear interactions (but we did not remove other predicted localizations from the automated call set).

As a companion to this curated interaction dataset, we also made use of the TAIR Interactome v 2.0 (Geisler-Lee, et al. 2007), which identifies proteins with direct physical interactions. We used all pairwise interactions to create a list of partners for each Araport11 protein (File S1). For organelle-targeted proteins, lists were further refined to include interacting partners that are targeted to the same subcellular compartment.

### Identification of direct cytonuclear contact sites within multisubunit enzyme complexes

In some cases, nuclear-encoded proteins may form part of a cytonuclear enzyme complex but still not physically contact a cytoplasmic gene product within the complex. Therefore, to identify direct cytonuclear interactions at the level of subunits and amino-acid residues, we mapped *A. thaliana* protein sequences to reference structures of 13 multisubunit enzyme complexes that are involved in OXPHOS, photosynthesis, protein translation, and fatty acid biosynthesis. Reference structures for these complexes were searched in the Protein Data Bank (PDB) and were chosen based on their completeness and relatedness to *A. thaliana* (File S1). We identified cytonuclear contact residues in these structures using the “find clashes/contacts” tool in Chimera version 1.12 (Pettersen, et al. 2004) with default contact settings except that VDW overlap was changed to ≥ − 1 to increase the sensitivity of detecting contacts. We determined homologous genes and residues in *A. thaliana* by querying the structural reference sequences with TAIR BLAST 2.2.8, and we aligned the resulting hits with MUSCLE as implemented in MEGA 7 (Kumar, et al. 2016) to identify the corresponding contact residues in *A. thaliana* genes.

## RESULTS AND DISCUSSION

CyMIRA is a detailed curation of *A. thaliana* cytonuclear interactions at the molecular level, which is available as supplementary material with this paper (File S1). Future updates will be disseminated via GitHub (https://github.com/dbsloan/cymira), and we have also generated a queryable web interface to extract specific subsets of the data: http://cymira.colostate.edu/.

Our initial automated predictions of organelle targeting based on nine existing databases (Table 1) identified a total of 4,130 nuclear-encoded protein-coding genes (1,256 mitochondrial-localized, 2,468 plastid-localized, and 406 dual-localized). The sampled databases differed greatly in their number of organelle-targeting predictions, and very few genes shared the same prediction across all nine databases (Figure 1). Because we limited our classification to predictions shared by at least two databases, there were thousands of genes that were excluded because they had a mitochondrial or plastid targeting prediction in only a single database (Figure 1). As such, taking the full union of predictions across all nine databases would have massively exceeded typical estimates of mitochondrial and plastid proteome content.

**Figure 1:**
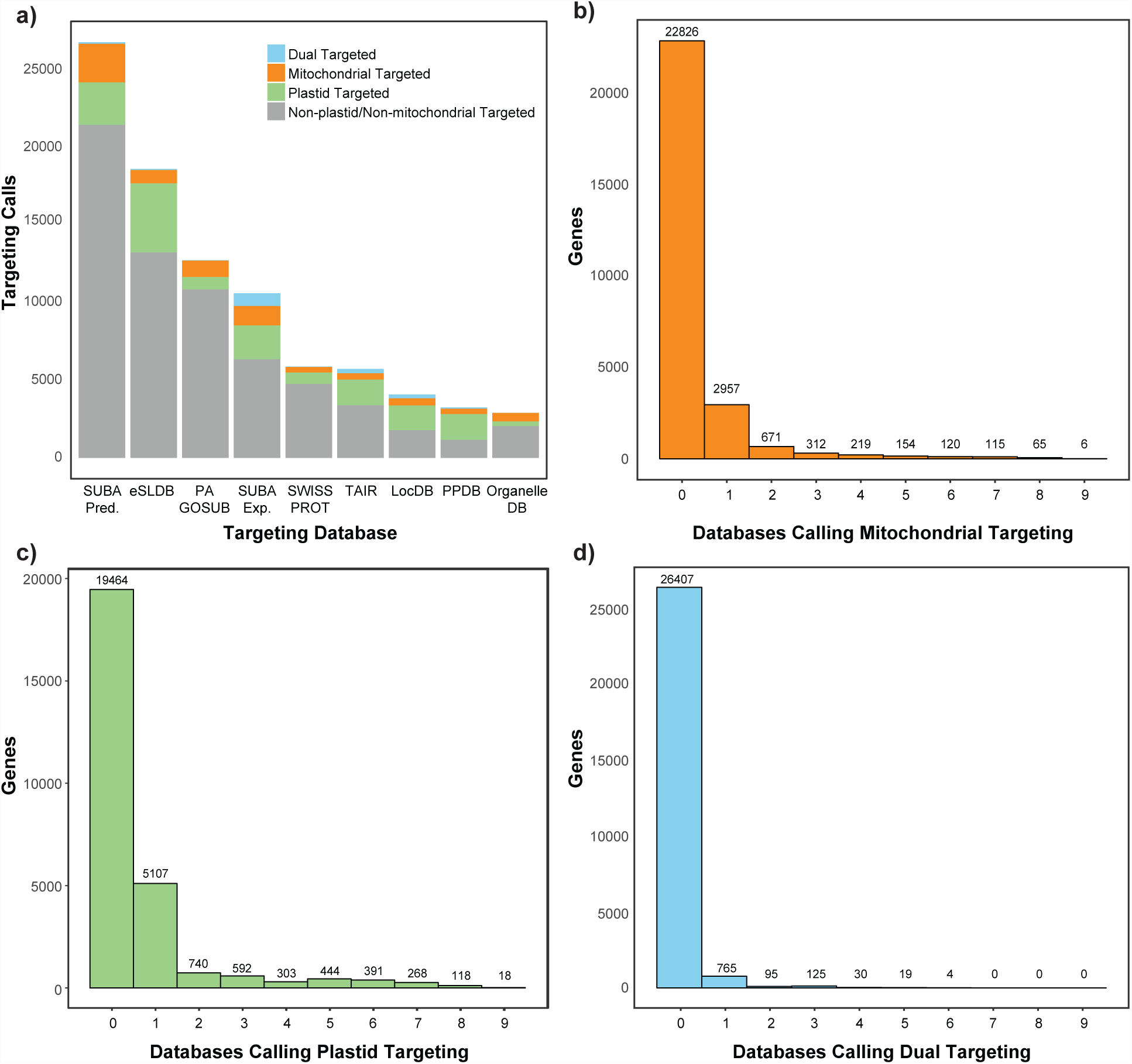
Summary of nine existing databases on subcellular protein targeting plants that were used to generate our automated targeting predictions.

Subsequent manual curation of proteins with direct cytonuclear interactions led to the inclusion of 138 new genes and changed the prediction for six genes that were initially identified as targeting just one organelle to dual targeting. As a result, our final organelle targeting count was 4,268 with 1,337 mitochondrial, 2,495 plastid, and 436 dual. Of these, 910 were classified as being involved in direct cytonuclear molecular interactions, meaning that they are components of chimeric cytonuclear enzyme complexes or directly interact with cytoplasmic DNA and/or RNA transcripts (Table 2). The majority of genes involved in these direct cytonuclear interactions were characterized as exclusively mitochondrial (535) or plastid (293), but there are also 82 dual targeted genes in this group, many of which are involved in DNA recombination/replication/repair, tRNA aminoacylation, and post-transcriptional RNA modifications (File S1).

Many studies have begun taking advantage of protein structural data to specifically investigate molecular evolution at the physical interface between contacting cytoplasmic and nuclear gene products (Osada and Akashi 2012; Havird, et al. 2015; Zhang, et al. 2015; Havird and McConie 2019; Yan, et al. 2019). We therefore used structural data from 13 protein complexes (Figure 2) to identify which nuclear subunits actually contact cytoplasmically encoded subunits within these complexes and their specific interacting amino acid positions (File S1). However, the efficacy of this structural mapping approach varied greatly depending on the completeness and phylogenetic relatedness of the reference structures. For many photosynthetic complexes, reference structures are available from angiosperms or even *A. thaliana* itself, but other complexes required use of structures from anciently divergent species, including bacteria and mammals (File S1), making inference of residue homology tenuous. Furthermore, even when structures from close relatives were available, they were sometimes known to be missing certain subunits (van Bezouwen, et al. 2017; Laughlin, et al. 2019). Therefore, we did not analyze many subunits within these complexes because of their absence from reference structures or low level of sequence similarity, designating them simply as not available (“NA”). Some additional subunits were classified only as “likely” or “not likely” to be involved in direct cytonuclear interactions because of low confidence in the reference mapping. Despite these limitations, structural data suggest that most nuclear-encoded proteins within these chimeric complexes do physically contact cytoplasmic gene products (91% of those for which assignments were made).

**Figure 2:**
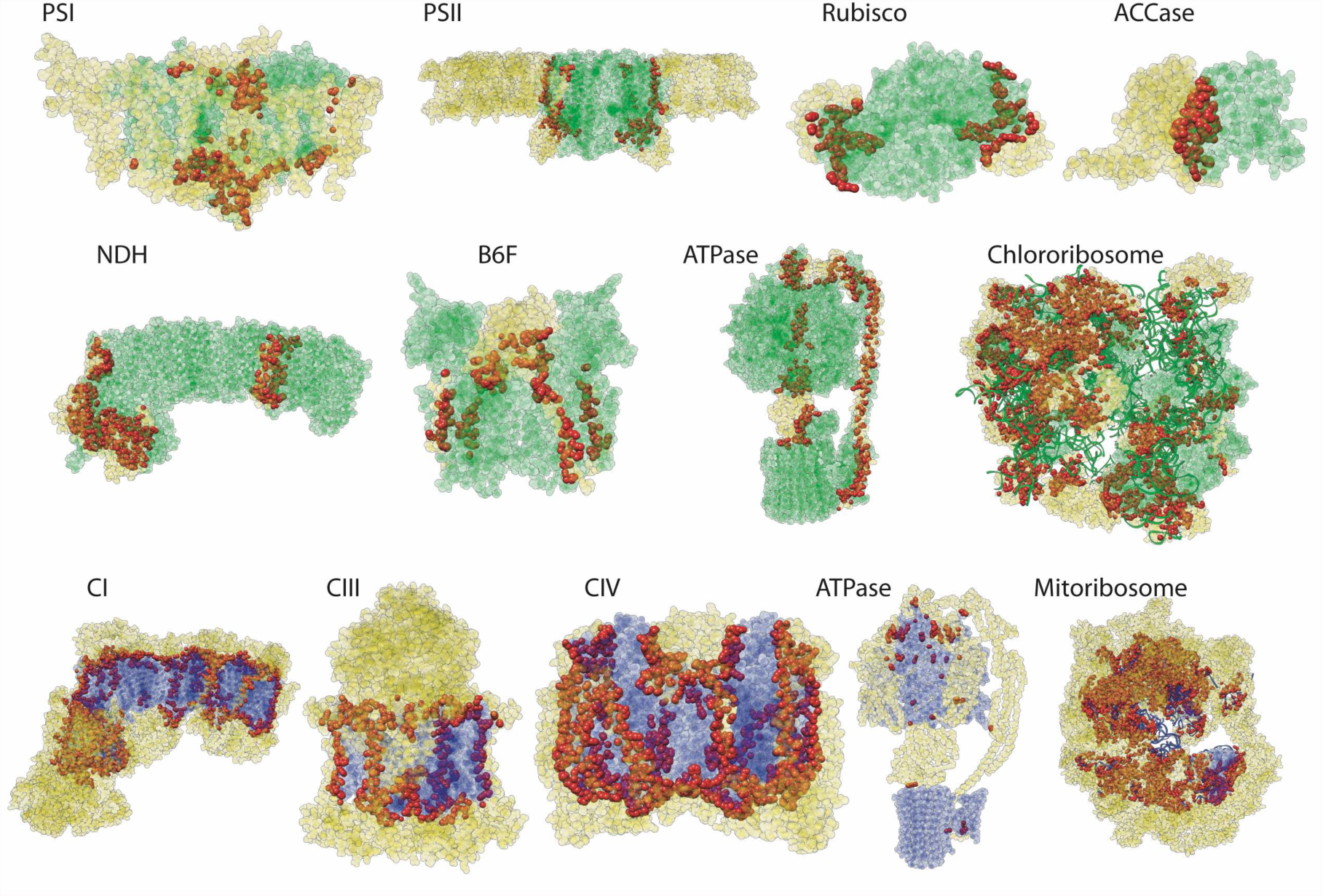
Chimeric cytonuclear protein complexes showing cytoplasmic-encoded, nuclear-encoded, and nuclear contact residues. Plastid-encoded residues are in green, mitochondrial-encoded residues are in purple, nuclear-encoded non-contact residues are in yellow, and nuclear-encoded contact residues are in red. Amino acids are shown as spheres, RNA is shown as ribbons. PDB accessions for reference structures: PSI: 2O01, PSII: 5MDX, rubisco: 5IU0, ACCase: 2F9Y, NDH: 6NBY, B6F: 1VF5, plastid ATPase: 6FKF, chlororibosome: 5MMM, CI: 5LNK, CIII: 1BGY, CIV: 1V54, mitochondrial ATPase: 5ARA, and mitoribosome: 3J9M.

Our goal in generating CyMIRA is to provide a standardized partitioning of plant nuclear gene content based on cytonuclear interactions at a molecular level to improve consistency across evolutionary genomic studies. One obvious need that will arise is to extend this *A. thaliana* annotation to genomic datasets from non-model plant species that lack the same level of functional data. Because of the extensive history of gene and whole-genome duplication and the associated process of neofunctionalization in plants (Panchy, et al. 2016), we recommend against relying solely on homology searches when porting the CyMIRA annotations to other species. Instead, we suggest combining such information with tools that perform *in silico* predictions of organelle targeting to increase confidence in assignments (Bannai, et al. 2002; Small, et al. 2004; Emanuelsson, et al. 2007; Sperschneider, et al. 2017).

A further complication in expanding to evolutionary studies across species is that the landscape of cytonuclear integration and interactions is rapidly shifting in plants. Unlike many eukaryotes in which the gene content in cytoplasmic genomes has reached a period of long-term stasis (Johnston and Williams 2016; Janouškovec, et al. 2017), flowering plants remain highly active in the process of endosymbiotic gene transfer to the nucleus (Timmis, et al. 2004). For example, our CyMIRA annotations do not include OXPHOS complex II because this is entirely nuclear-encoded in *A. thaliana*. In contrast, many other angiosperms have retained functional complex II genes (*sdh3* and/or *sdh4*) in their mitochondrial genomes. Ribosomal subunits are also subject to ongoing functional transfers to the nucleus, resulting in substantial heterogeneity in cytoplasmic gene content across angiosperms (Adams, et al. 2002). Therefore, species-specific additions and deletions to this dataset, even at the whole complex level, should be considered based on the retained cytoplasmic gene content in each lineage. Although this continued need for refinement across phylogenetic scales undoubtedly poses a challenge for future studies, the dynamic nature of cytoplasmic genomes in plants is also one of the strongest motivations for studying cytonuclear interactions in these systems.

In summary, the proliferation of plant genomic resources makes this an exciting time to take studies of cytonuclear biology to a genome-wide level, and methodological consistency will be key to the efficacy of such efforts. We hope that CyMIRA will serve as useful community resource in this respect.

## ACKNOWLEDGEMENTS

We thank Stephane Bentolila, Hans-Peter Braun, José Gualberto, Oren Ostersetzer-Biran, Mareike Schallenberg-Rüdinger, and Ian Small for helpful discussion on different types of cytonuclear molecular interactions. This work was supported by NSF grants IOS-1829176 and MCB-1733227, an NSF GAUSSI graduate research fellowship (DGE-1450032), and start-up funds from the University of Texas.

---

**Table S1:**
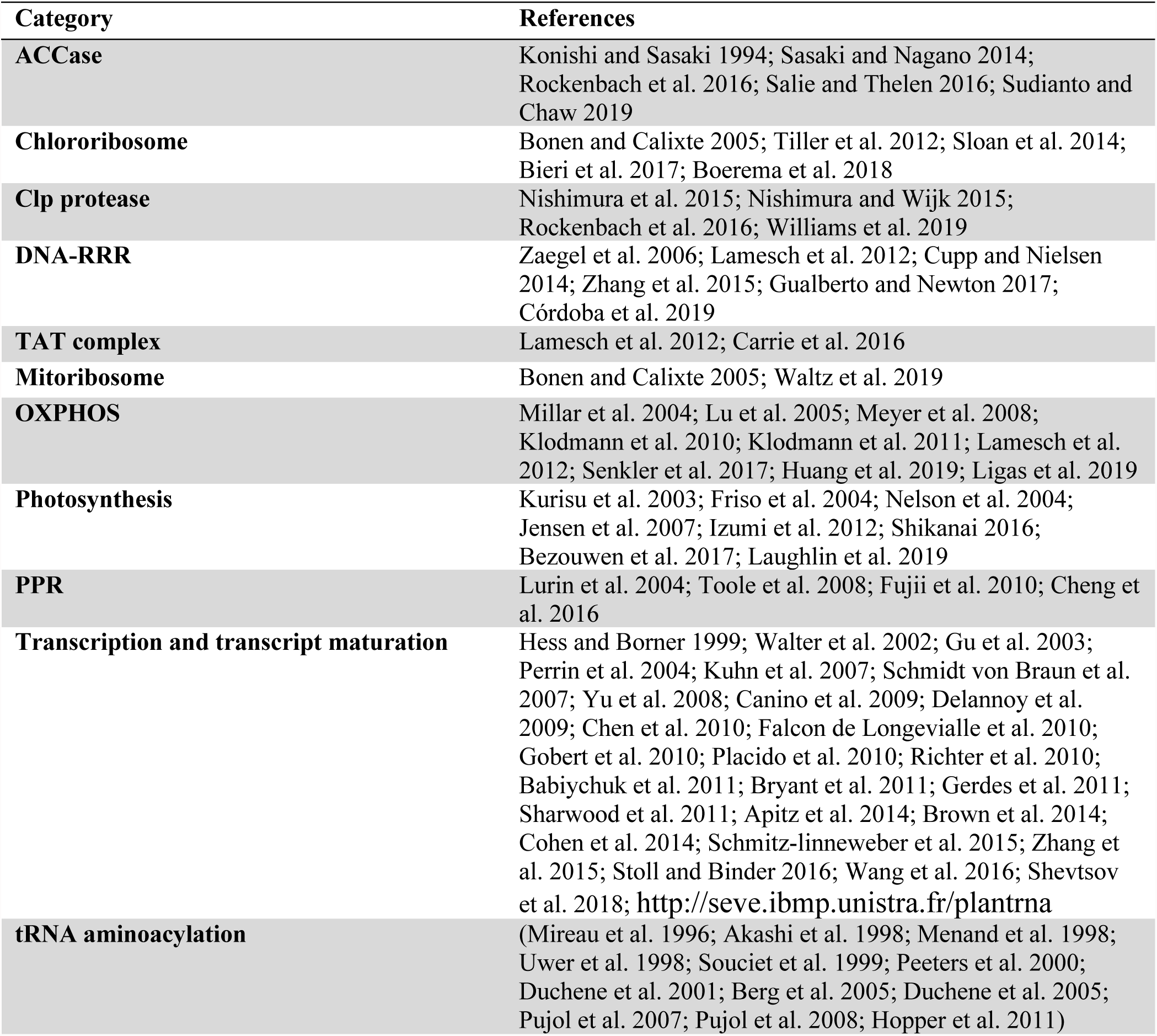
Full list of references used for manual curation of cytonuclear complexes

